# *In vivo* RNA targeting of point mutations via suppressor tRNAs and adenosine deaminases

**DOI:** 10.1101/210278

**Authors:** Dhruva Katrekar, Prashant Mali

## Abstract

Point mutations underlie many genetic diseases. In this regard, while programmable DNA nucleases have been used to repair mutations, their use for gene therapy poses multiple challenges: one, efficiency of homologous recombination is typically low in cells; two, an active nuclease presents a risk of introducing permanent off-target mutations; and three, prevalent programmable nucleases typically comprise elements of non-human origin raising the potential of *in vivo* immunogenicity. In light of these, approaches to instead directly target RNA, and use of molecular machinery native to the host would be highly desirable. Towards this, we engineered and optimized two complementary approaches, referred together hereon as **tRiAD**, based on the use of **tR**NAs **i**n codon suppression and **a**denosine **d**eaminases in RNA editing. Specifically, by delivering modified endogenous tRNAs and/or the RNA editing enzyme ADAR2 and an associated guiding RNA (adRNA) via adeno-associated viruses, we enabled premature stop codon read-through and correction in the *mdx* mouse model of muscular dystrophy that harbors a nonsense mutation in the dystrophin gene. We further demonstrated inducible restoration of dystrophin expression by pyrolysyl-tRNA mediated incorporation of unnatural amino acids (UAAs) at the stop codon. Additionally, we also engineered ADAR2 mediated correction of a point mutation in liver RNA of the *spf*^*ash*^ mouse model of ornithine transcarbamylase (OTC) deficiency. Taken together, our results establish the use of suppressor tRNAs and ADAR2 for *in vivo* RNA targeting, and this integrated tRiAD approach is robust, genomically scarless, and potentially non-immunogenic as it utilizes effector RNAs and human proteins.

## MAIN

Genome engineering methodologies coupled with rapidly advancing synthetic biology toolsets are enabling powerful capabilities to perturb genomes for deciphering function, programming novel function, and repairing aberrant function. In particular, programmable DNA nucleases, such as meganucleases, zinc finger nucleases (ZFNs), transcription activator-like effector nucleases (TALENs), and CRISPR-Cas, have been widely used to engineer genomes^1–5^. Their use in gene therapy however poses at least three major challenges: one, the efficiency of homologous recombination versus non-homologous end joining is typically low, particularly in post-mitotic cells that comprise the vast majority of the adult body^6,7^; two, an active nuclease always poses the threat of introducing permanent off-target mutations, thus presenting formidable challenges in both engineering exquisite nuclease specificity without compromising activity, as well as in tight regulation of the nuclease dose and duration in target cells^8,9^; and three, prevalent programmable nucleases are of prokaryotic origin or bear domains that are of non-human origin raising a significant risk of immunogenicity in *in vivo* therapeutic applications^10,11^. For genomic mutations that lead to alteration in protein function, which represent a significant majority of disease causing mutations, approaches to instead directly target RNA would be highly desirable. Leveraging the aspect that RNA is generally more accessible to oligonucleotide mediated targeting without the need for enabling proteins than double-stranded DNA, and building on the advances in tRNA mediated codon suppression^12–15^ and genetic code expansion^16–18^, as well as adenosine deaminase mediated RNA editing^19–23^, we engineer and optimize here an integrated platform for RNA targeting, and demonstrate its efficacy in *in vitro* and *in vivo* applications.

We focused first on establishing the system for targeting nonsense mutations. Nearly one-third of the alleles responsible for causing genetic disorders carry premature stop codons, leading to the production of mRNA with truncated reading frames^24^. Nonsense mutations are also highly prevalent in cancers, typically occurring in tumor suppressor genes^25–27^. For instance, about 8% of all mutations in p53 are nonsense mutations^28^. Overall, nonsense mutations are responsible for 11% of all described gene lesions causing inheritable human disease and close to 20% of disease-associated single base substitutions that affect the coding regions of genes^29,30^. Commonly used strategies to target nonsense mutations that have shown promising results *in vivo,* include the use of aminoglycosides and PTC124 which promote stop codon read-through^28,31–36^, exon skipping therapies^37–40^, and more recently nuclease based DNA editing approaches^41–46^. Aminoglycosides are mainly used as antibacterial drugs that act by targeting the 16S rRNA subunit of the bacterial ribosome, leading to the inhibition of protein synthesis in prokaryotes^47,48^. However, these drugs are less efficient in eukaryotes and enable read-through of stop codons, resulting in the formation of both full length and truncated proteins^49^. Drugs such as gentamycin have however shown mixed results in clinical trials for premature stop codon read-through in Duchenne Muscular Dystrophy and Cystic Fibrosis with variable efficiencies^33,50,51^. Exon skipping strategies in turn find clinical application only in specific cases where truncated proteins retain wild type functionality^37^. On the contrary, genomic targeting approaches are more promising as they involve the direct modulation of the expression levels or the direct correction of aberrant genes. For instance, in order to restore expression of a gene bearing a premature stop codon, gene editing based approaches involving introduction of a functional gene to replace it^52,53^, or alternatively, in frame excision of the region bearing the premature stop codon have indeed shown encouraging results *in vitro* and *in vivo*^42–46^. However, as outlined earlier, key issues associated with the current low efficiency of homology directed repair (HDR), possibility of permanent off targeting, and potential to elicit an immune response, remain to be overcome.

In light of the above, we explored two independent but complementary approaches to directly target RNA. First, we focused on engineering robust nonsense codon suppression via suppressor tRNAs. The anticodon stem of human serine^12^, leucine^13^, and arginine^14^ tRNAs have been previously mutated to generate suppressor tRNAs. These suppressor tRNAs can be charged by the endogenous aminoacyl tRNA synthetases (aaRS) and have been shown to bring about partial restoration of expression of proteins encoded by mRNA harboring the nonsense mutations in xeroderma pigmentosum cells^14^. Additionally, orthogonal suppressor tRNA-aaRS pairs have widely been used to incorporate unnatural amino acids (UAAs) into a protein leading to an expansion of the genetic code^16^. Although the use of suppressor tRNAs for premature stop codon read-through of endogenous non-sense mutations has been attempted *in vivo* in mice, these prior studies relied only on plasmid delivery and the use of robust and optimized delivery formats was not explored^54,55^. Additionally, the potential use of UAA based inducible *in vivo* suppression of a disease-causing endogenous nonsense mutation has not been explored either. Based on the above, we first modified the anticodon stems of serine, arginine and leucine tRNAs to create suppressor tRNAs^12–14^ targeting all three stop codons and evaluated these side-by-side in cells using GFP reporters harboring corresponding nonsense mutations replacing the Y39 residue. Among these, the serine suppressor tRNA demonstrated the most consistent and robust results (**Figure 1a, Supplementary Figure 1a**). To also engineer inducible codon suppression, we next utilized the pyrolysyl-tRNA/aminoacyl tRNA synthetase (aaRS) pair from *Methanosarcina barkeri* (MbPylRS)^56,57^ and cloned it into AAV vectors. This enabled incorporation of UAAs at a stop codon. Notably, we found that adding a second copy of the tRNA into the expression vector significantly boosted suppression efficiencies (**Figure 1b**). We further systematically evaluated additional aminoacyl tRNA synthetases from *Methanosarcina mazei* (MmPylRS)^58^ and an *N*ɛ-acetyl-lysyl-tRNA synthetase (AcKRS)^18^, and also explored varying the number of tRNAs copies per vector to up to four (**Supplementary Figure 1b**).

**Figure 1:**
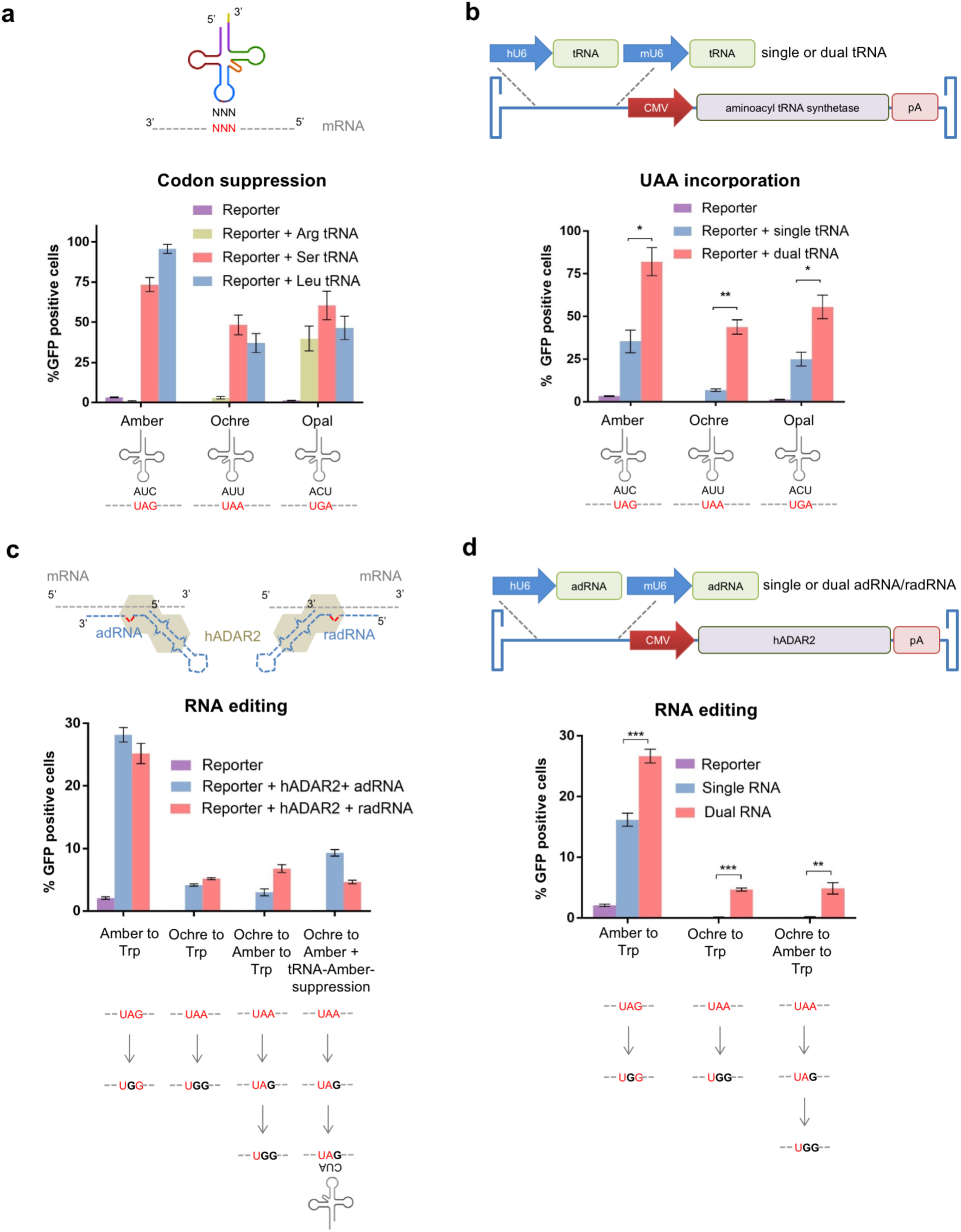
*In vitro* suppression and editing of stop codons in GFP reporter mRNA. **(a)** Arginine, serine and leucine suppressor tRNAs targeting amber, ochre and opal stop codons. **(b)** Orthogonal tRNA/aaRS (MbPylRS) based suppression of amber, ochre and opal stop codons in the presence of one or two copies of the pyrolysyl-tRNA delivered via an AAV vector and 1mM Nɛ-Boc-L-Lysine. **(c)** ADAR2 based RNA editing efficiencies of amber and ochre stop codons, in one-step, two-steps, or in combination with suppressor tRNAs. **(d)** ADAR2 based RNA editing efficiencies of amber and ochre stop codons in the presence of one or two copies of the adRNA, delivered via an AAV vector.

Concurrently we also engineered a system for sequence-specific targeted RNA editing via adenosine deaminase enzymes. Adenosine to Inosine (A to I) editing is a common post-transcriptional modification in RNA, catalyzed by adenosine deaminases acting on RNA (ADARs)^19^. Inosine is a deaminated form of adenosine and is biochemically recognized as guanine. Recently, multiple studies have demonstrated the engineering of ADAR2 mediated targeting *in vitro*^21–23^, and a study also demonstrated correction of the nonsense mutation in CFTR in xenopus oocytes^21^. Building on this, we engineered here a robust system for sequence-specific targeted RNA editing *in vitro* and *in vivo*, utilizing the human ADAR2 enzyme and an associated ADAR2 guide RNA (adRNAs) engineered from its naturally occurring substrate GluR2 pre-mRNA^21–23^. This ADAR2 guiding RNA comprises an ADAR-recruiting domain and a programmable antisense region that is complementary to a specified target RNA sequence. We first evaluated the RNA editing efficiency of this system *in vitro* by co-transfecting our constructs with GFP reporters harboring a non-sense amber or ochre mutation at Y39. Specifically, we utilized two editing approaches to engineer the editing of both adenosines in the ochre stop codon: a one-step mechanism where both the adenosines are edited simultaneously or a two-step mechanism wherein editing takes place sequentially. In addition, we also explored the possibility of conversion of an ochre codon to an amber codon followed by amber suppression to restore GFP expression. All three approaches enabled robust restoration of GFP expression (**Figure 1c, Supplementary Figure 2a**). We next constructed AAV vectors delivering an adRNA or reverse oriented adRNA (radRNA) along with the ADAR2 enzyme. Similar to tRNA mediated codon suppression, addition of a second copy of the adRNA/radRNA also significantly improved the targeting efficiency (**Figure 1d**). We further systematically evaluated modified ADAR recruiting domains, and a range of RNA targeting antisense designs of varying lengths and the number of nucleotides intervening the target A and the R/G motif of the adRNA^22^ (**Supplementary Figure 2b**), yielding a panel of efficient adRNA designs (**Supplementary Figure 2c**).

**Figure 2:**
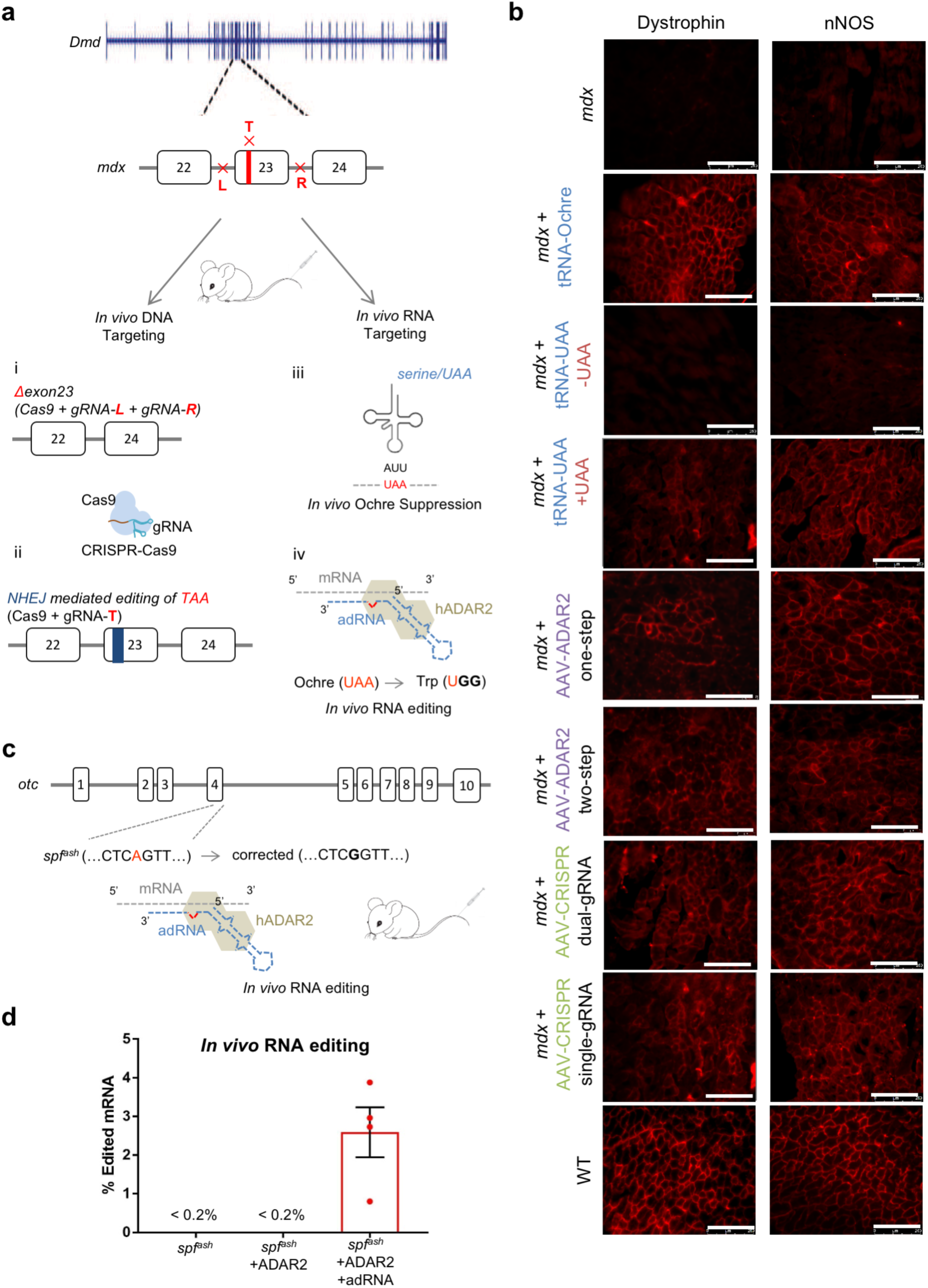
*In vivo* RNA targeting in mouse models of human disease. **(a)** Schematic of the DNA and RNA targeting approaches to restore dystrophin expression in *mdx* mice. DNA targeting is enabled by the CRISPR-Cas9 system, using either (i) a dual gRNA based approach leading to the excision of exon 23 or (ii) a single gRNA based approach that relies on NHEJ to restore the full-length dystrophin. RNA targeting is enabled either via (iii) tRNA suppression or (iv) ADAR2 based editing of the ochre stop codon. **(b)** Immunofluorescence staining for dystrophin and nNOS in controls and treated samples (scale bars: 250μm). **(c)** Schematic of the OTC locus in *spf*^*ash*^ mice which have a G->A point mutation at a donor splice site in the last nucleotide of exon 4, and approach for correction of mutant OTC mRNA via ADAR2 mediated RNA editing **(d)** *In vivo* A->G RNA editing efficiencies in corresponding treated *spf*^*ash*^ mice.

Based on the above *in vitro* optimizations, we next tested our system for *in vivo* RNA targeting. We focused first on the *mdx* mouse model for Duchenne muscular dystrophy (DMD)^59^ which bears an ochre stop site in exon 23 of the dystrophin gene^60,61^. Recent studies utilizing the CRISPR-Cas9 system have shown promising results in the prevention^41^ and partial functional recovery^42–45^ of DMD by making changes in exon 23 at the DNA level in the *mdx* mouse. We thus concurrently evaluated four approaches (**Figure 2a**): one, modified endogenous tRNAs targeting the stop codon of interest; two, pyrolysyl-tRNA/aaRS pairs that can incorporate UAAs at the premature stop codon; three, ADAR2 based correction of the nonsense mutation; and, four, CRISPR-Cas9 based genome targeting to benchmark the above results. The use of modified endogenous tRNAs for targeting a stop codon has the advantage that these tRNA can be charged by endogenous aminoacyl-tRNA synthetases using an amino acid already present in the organism and are thus unlikely to produce an immune response. The use of pyrolysyl-tRNAs in turn makes the system inducible on-demand in the presence of the corresponding UAA. Finally, ADAR2 based RNA editing is site-specific, and due to the enzyme’s human origin, is less likely to elicit an immune response.

Specifically, as serine suppressor tRNAs gave us the best results *in vitro*, we designed an AAV carrying two copies of the serine suppressor tRNA targeting the ochre stop codon. Correspondingly, the tibalis anterior (TA) or gastrocnemius of *mdx* mice were injected with AAV8-dual-ser-ochre-tRNA. Mice muscle were harvested after 2, 4, and 8 weeks. Progressively improved restoration of dystrophin expression was seen over time, with the mice harvested after 8 weeks showing the highest degree of restoration (**Figure 2b**, **Supplementary Figure 3a**). In addition, neuronal nitric oxide synthase (nNOS) activity was restored at the sarcolemma which is absent in *mdx* mice due to the absence of the nNOS binding site in the dystrophin protein (**Figure 2b**). We also demonstrated the inducible restoration of dystrophin expression in the presence of UAAs. Specifically, a vector carrying two copies of the pyrolysyl-tRNA targeting the ochre stop codon and MbPylRS was constructed and injected into the TA or gastrocnemius of *mdx* mice, and the mice were divided into two groups: one that was administered the UAA and a control group that was not. As expected, only mice that were provided the UAA showed a restoration of dystrophin expression (**Figure 2b**).

We next evaluated our ADAR2 mediated site-specific RNA editing approach in this mouse model. To test the efficiency of our system in editing both adenosines in the ochre stop codon in *mdx* DMD mRNA, we first optimized our constructs *in vitro* with a reporter plasmid bearing a fragment of the *mdx* DMD mRNA in HEK293T cells. Sanger sequencing and NGS analysis confirmed successful targeting (**Supplementary Figure 3b**). We next packaged the optimized constructs into AAV8, and similar to the suppressor tRNA studies injected the tibialis anterior (TA) or gastrocnemius of *mdx* mice. Eight weeks post injection, TA and gastrocnemius muscles were collected from *mdx* mice, wild type mice, and mice treated with adRNA targeting and non-targeting controls. IHC revealed clear restoration of dystrophin expression (**Figure 2b**). In addition, nNOS activity was also restored at the sarcolemma (**Figure 2b**).

Furthermore, to benchmark the tRiAD approach we also targeted the *mdx* mice via CRISPR based genome editing of the nonsense mutation. We injected vectors bearing dual-gRNAs to excise exon 23, and also a single guide RNA directly targeting the premature stop codon, allowing for a chance NHEJ mediated in frame disruption of the non-sense mutation. Expectedly, both approaches led to restoration of dystrophin expression in a subset of the muscle cells (**Figure 2b**).

Finally, we also evaluated the ADAR2 mediated RNA targeting in an independent mouse model of human disease. Specifically, we focused on the male sparse fur ash (*spf*^*ash*^) mouse model of ornithine transcarbamylase (OTC) deficiency. The *spf*^*ash*^ mice have a G->A point mutation in the last nucleotide of the fourth exon of the OTC gene, which leads to OTC mRNA deficiency and production of mutant protein^62^. Recent studies have demonstrated the use of CRISPR-Cas9 and homologous recombination based strategies for robust correction of the mutation in neonatal mice^63^. However, gene correction via homology-directed repair (HDR) in adult mice was inefficient and led to genomic deletions which proved to be lethal as they compromised liver function in targeted mice^63^. To test the effectiveness of our system in editing the point mutation in *spf*^*ash*^ OTC mRNA (**Figure 2c**), we first evaluated our constructs *in vitro* with a plasmid bearing a fragment of the *spf*^*ash*^ OTC mRNA in HEK293T cells. Sanger sequencing and next generation sequencing (NGS) analysis confirmed robust RNA editing efficiencies (**Supplementary Figure 3c**). We then packaged our constructs into AAV8, which has high liver tropism^63^, and injected 10-12 week old *spf*^*ash*^ mice. Four weeks post injection, we collected liver samples from *spf^ash^,* wild-type litter mates, and *spf*^*ash*^ mice treated with the ADAR2 targeting and non-targeting vectors and evaluated targeting efficiency via NGS. Notably, significant RNA editing rates in the range of 0.85-4% were observed in treated mice (**Figure 2d**, **Supplementary Figure 3d**), further confirming the utility of this approach in *in vivo* editing of endogenous RNA.

Taken together, our results establish the use of suppressor tRNAs and ADAR2 as potential strategies for *in vivo* RNA targeting of point mutations. Specifically, by optimizing delivery, we first demonstrated robust and inducible stop codon read-through via the use of suppressor tRNAs. The delivery of modified endogenous suppressor tRNAs for premature stop read-through has several potential advantages: it lacks the toxicity associated with read-through drugs such as gentamycin and can be used to bring about efficient stop codon read-through in post-mitotic cells. In addition, being of endogenous origin, it is not likely to elicit a strong immune response. The inducibility enabled by the UAA based systems, albeit non-native, could further provide tight regulation over the expression of genes. We however note too that an important caveat to this overall strategy, analogous to the read-through drugs, is that suppressor tRNA based approaches will lead to the read-through of other non-target stop codons. In this regard, we also demonstrated ADAR2 based site-specific correction of point mutations in RNA. We note that potential off-targets in RNA are limited as compared to DNA as the transcriptome is only a small subset of the genome. Secondly, even if off-targets exist, the presence of an A within the target window is required for the enzyme to create an off-target A->G change. Lastly, the off-target effects will be transient. Thus, overall off-target effects due to a RNA editing enzyme such as ADAR2 are likely to be limited, although enzyme processivity, promiscuity, and off-target hybridization of the antisense domain of the adRNA needs to be studied thoroughly. ADAR2 being of human origin is also less likely to elicit an immune response, while enabling more site-specific editing of RNA compared to the suppressor tRNA approach.

We note that compared to the tRiAD based RNA targeting approaches above, CRISPR based genome targeting approaches currently show faster kinetics and greater degree of mutant protein restoration. We however anticipate that systematic engineering and directed evolution of the ADAR2 could help improve the editing efficiency and also eliminate the intrinsic biases of the ADAR2 for certain sequences^64^. The demonstration of site-specific A->G mRNA editing *in vivo* also opens up the door to future site-specific C->T editing via targeted recruitment of cytosine deaminases, thereby potentially expanding the repertoire of RNA editing tools. However, an important consideration while targeting RNA for gene therapy via the use of non-integrating vectors such as AAVs, is the necessity for periodic re-administration of the effector constructs due to the typically limited half-life of edited mRNAs. Secondly, RNA folding, post translational modifications, and RNA binding proteins might also impact accessibility of target sites in the RNA. Chemical and structural modifications in tRNAs and adRNAs^65,66^, or co-expression of shielding proteins might help improve RNA stability and specificity, and improve the efficacy of the above approaches. With progressive improvements, we thus anticipate this integrated tRiAD toolset will have broad implications in both applied life sciences as well as fundamental research.

## ACKNOWLEDGEMENTS

We thank Genghao Chen, Atharv Worlikar, Ana Moreno, Dongxin Zhao, Udit Parekh, and other members of the Mali lab for advice and help with experiments, Yu-Ru Shih and Shyni Varghese for their advice and help on assays for the DMD experiments, and the Salk GT3 viral core for help with AAV production. This work was supported by UCSD Institutional Funds, the Burroughs Wellcome Fund (1013926), the March of Dimes Foundation (5-FY15-450), the Kimmel Foundation (SKF-16-150), and NIH grant (R01HG009285). D.K. and P.M. have filed patents based on this work.

## METHODS

### Vector design and construction

To construct the GFP reporters – GFP-Amber, GFP-Ochre and GFP-Opal, three gene blocks were synthesized with ‘TAG’, ‘TAA’ and ‘TGA’ respectively replacing the Y39 residue of the wild type GFP and were cloned downstream of a CAG promoter. One or two copies of the endogenous suppressor tRNAs were cloned into an AAV vector containing a human U6 and mouse U6 promoter. Pyrolysyl tRNAs and adRNAs/radRNAs were cloned into an AAV vector containing a human U6 and mouse U6 promoter along with a CMV promoter driving the expression of MbPylRS/MmPylRS/AcKRS or hADAR2 respectively.

### Mammalian cell culture and transfection

All HEK 293T cells were grown in Dulbecco’s Modified Eagle Medium supplemented with 10% FBS and 1% Antibiotic-Antimycotic (Thermo Fisher) in an incubator at 37 °C and 5% CO2 atmosphere. All *in vitro* transfection experiments were carried out in HEK 293T cells using the commercial transfection reagent Lipofectamine 2000 (Thermo Fisher). All *in vitro* suppression and editing experiments were carried out in 24 well plates using 500ng of reporter plasmid and 1000ng of the suppressor tRNA/aaRS plasmid or the adRNA/ADAR2 plasmid. Cells were transfected at 30% confluence. Cells were harvested 48 and 72 hours post transfection for quantification of suppression and editing respectively. The UAAs Nɛ-Boc-L-Lysine (Chemimpex) and Nɛ-Acetyl-L-Lysine (Sigma) were added to the media at the desired concentration before transfection.

### Production of AAV vectors

Virus was prepared using the protocol from the Gene Transfer, Targeting and Therapeutics (GT3) core at the Salk Institute of Biological Studies (La Jolla, CA). AAV8 particles were produced using HEK 293T cells via the triple transfection method and purified via an iodixanol gradient^67^. Confluency at transfection was about 80%. Two hours prior to transfection, DMEM supplemented with 10% FBS was added to the HEK 293T cells. Each virus was produced in 5 × 15 cm plates, where each plate was transfected with 7.5 ug of pXR-8, 7.5 of ug recombinant transfer vector, 7.5 ug of pHelper vector using PEI (1ug/uL linear PEI in 1xDPBS pH 4.5, using HCl) at a PEI:DNA mass ratio of 4:1. The mixture was incubated for 10 minutes at RT and then applied dropwise onto the cell media. The virus was harvested after 72 hours and purified using an iodixanol density gradient ultracentrifugation method. The virus was then dialyzed with 1 x PBS (pH 7.2) supplemented with 50 mM NaCl and 0.0001% of Pluronic F68 (Thermo Fisher) using 50kDA filters (Millipore), to a final volume of ~1 mL and quantified by qPCR using primers specific to the ITR region, against a standard (ATCC VR-1616).

*AAV-ITR-F: 5’-CGGCCTCAGTGAGCGA-3’ and AAV-ITR-R: 5’-GGAACCCCTAGTGATGGAGTT-3’*.

### RNA isolation and Next Generation Sequencing library preparation

RNA from gastrocnemius or TA muscles of *mdx* mice or livers of *spf*^*ash*^ mice was extracted using the RNeasy Plus Universal Mini Kit (Qiagen), according to the manufacturer’s protocol. Next generation sequencing libraries were prepared as follows. cDNA was synthesized using the Protoscript II First Strand cDNA synthesis Kit (New England Biolabs). Briefly, 500 ng of input cDNA was amplified by PCR with primers that amplify 150 bp surrounding the sites of interest using KAPA Hifi HotStart PCR Mix (Kapa Biosystems). PCR products were gel purified (Qiagen Gel Extraction kit), and further purified (Qiagen PCR Purification Kit) to eliminate byproducts. Library construction was done with NEBNext Multiplex Oligos for Illumina kit (NEB). 10 ng of input DNA was amplified with indexing primers. Samples were then pooled and loaded on an Illumina Miseq (150 single-end run) at 5nM concentrations. Data analysis was performed using CRISPResso^68^.

### Animal experiments

#### AAV Injections

All animal procedures were performed in accordance with protocols approved by the Institutional Animal Care and Use Committee (IACUC) of the University of California, San Diego. All mice were acquired from Jackson labs. AAVs were injected into the gastrocnemius or TA muscle of *mdx* mice (6-10 weeks old) using 2.5E+12 vg/muscle. AAVs were injected into *spf*^*ash*^ mice via retro-orbital injections using 3E+12 vg/mouse.

#### UAA administration

Mice were fed water containing 20 mg/ml Nɛ-Boc-L-Lysine (Chemimpex) for one month. Mice were also administered the 30 mg Nɛ-Boc-L-Lysine via IP injections, thrice a week.

### Immunofluorescence

Harvested gastrocnemius or TA muscles were placed in molds containing OCT compound (VWR) and flash frozen in liquid nitrogen. 20 μm sections were cut onto pre-treated histological slides. Slides were fixed using 4% Paraformaldehyde. Dystrophin was detected with a rabbit polyclonal antibody against the N-terminal domain of dystrophin (1:100, Abcam 15277) followed by a donkey anti-rabbit Alexa 546 secondary antibody (1:250, Thermo Fisher).

---

**Supplementary Figure 1:**
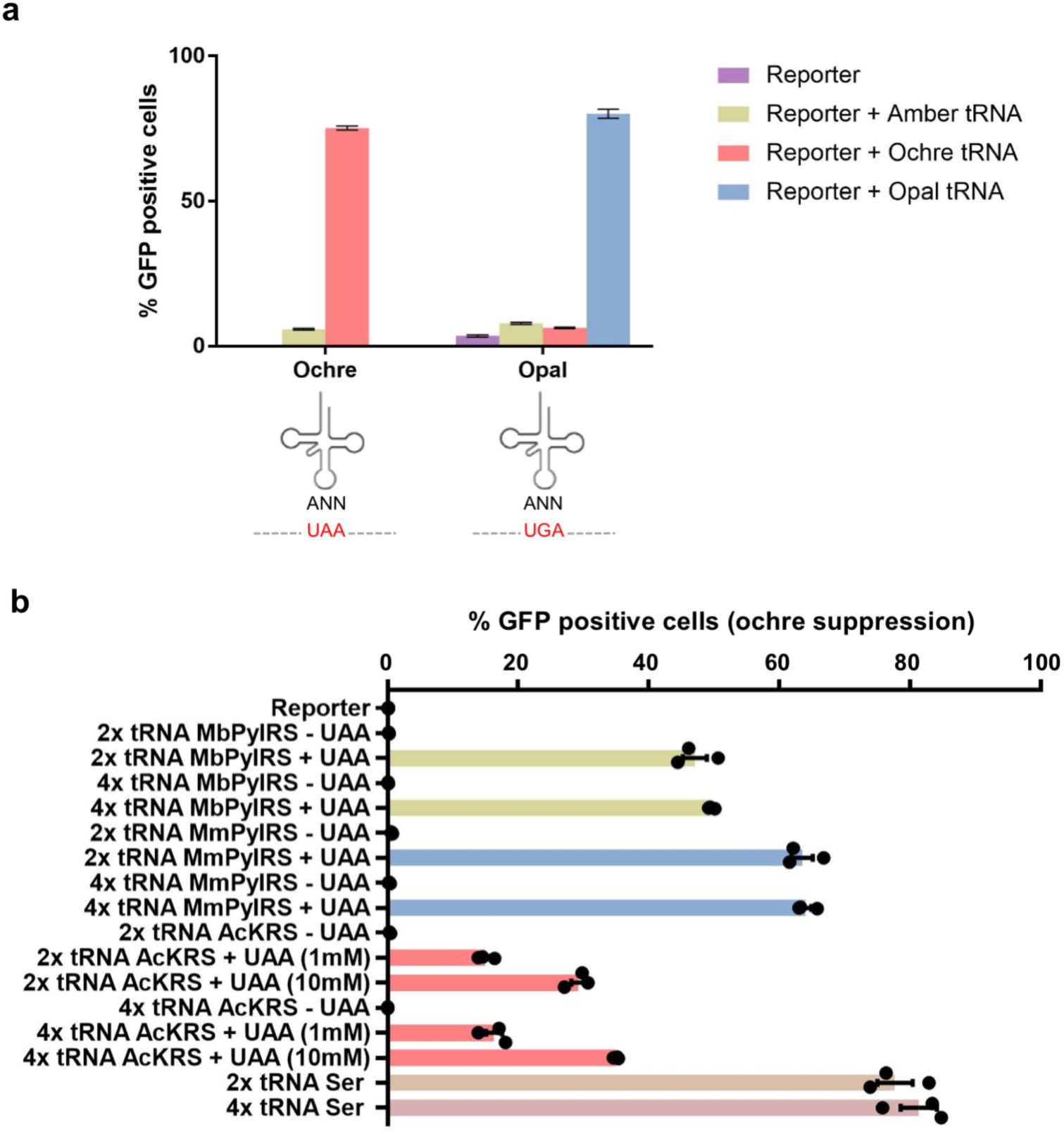
*In vitro* tRNA suppression evaluation and optimization. **(a)** Specificity of modified serine suppressor tRNAs for ochre and opal stop codons. **(b)** Ochre stop codon suppression efficiency utilizing three different aaRS: MbPylRS, MmPylRS and AcKRS, and two or four copies of the pyroysyl-tRNA, or serine suppressor tRNA, all delivered using an AAV vector. MbPylRS, MmPylRS: 1mM Nɛ-Boc-L-Lysine; AcKRS: 1 or 10mM Nɛ-Acetyl-L- Lysine.

**Supplementary Figure 2:**
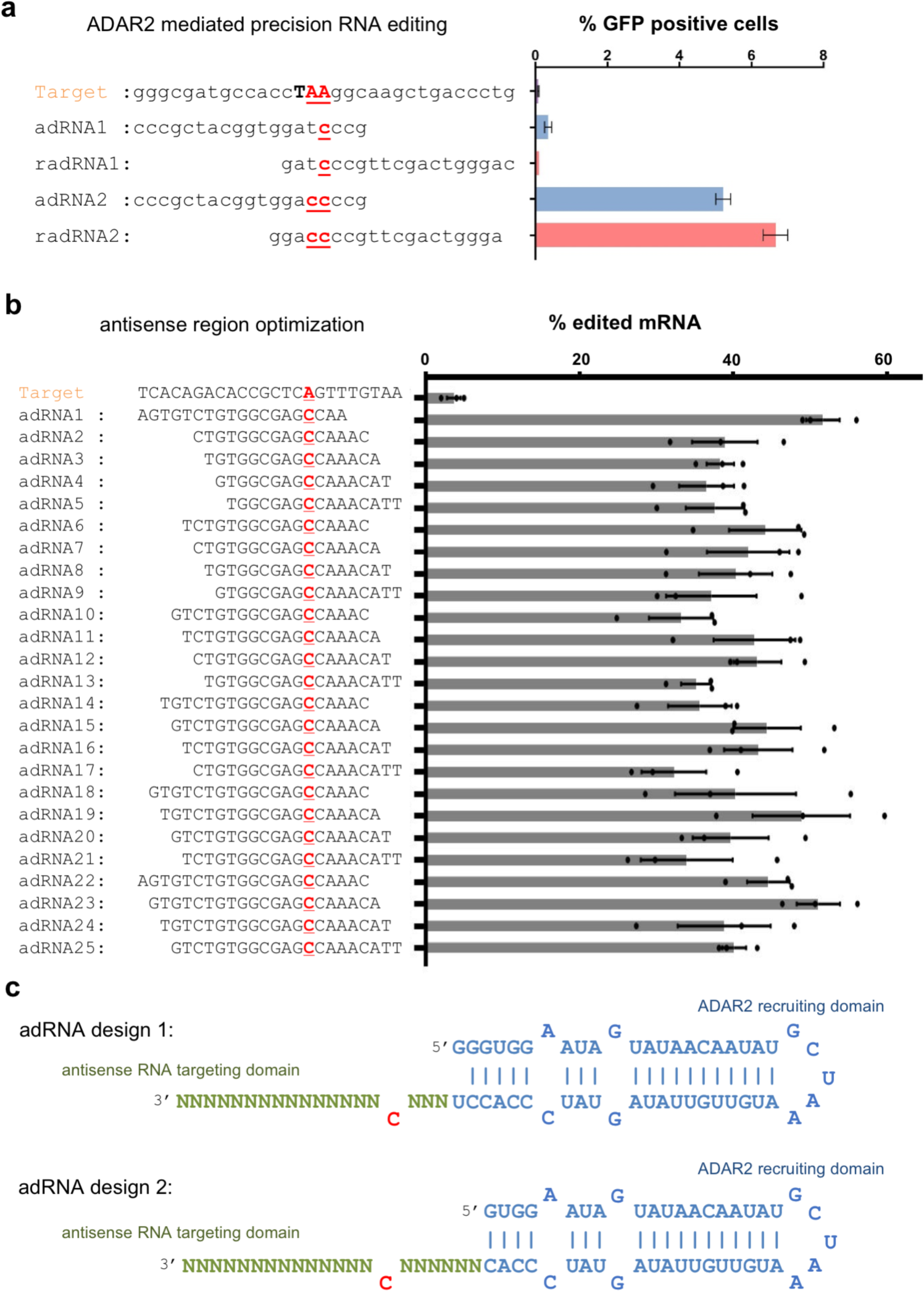
*In vitro* ADAR2 mediated site-specific RNA editing evaluation and optimization. **(a)** GFP expression is restored when adRNA/radRNA has two mismatches corresponding to the two adenosines in the ochre stop codon. Presence of a single mismatch results in the formation of an amber or opal stop codon. **(b)** Optimization of adRNA antisense region: length and distance from the ADAR2 recruiting region were systematically varied, and editing efficiency calculated as a ratio of Sanger peak heights G/(A+G) (n=3 independent transfections). **(c)** Panel of optimized adRNA designs.

**Supplementary Figure 3:**
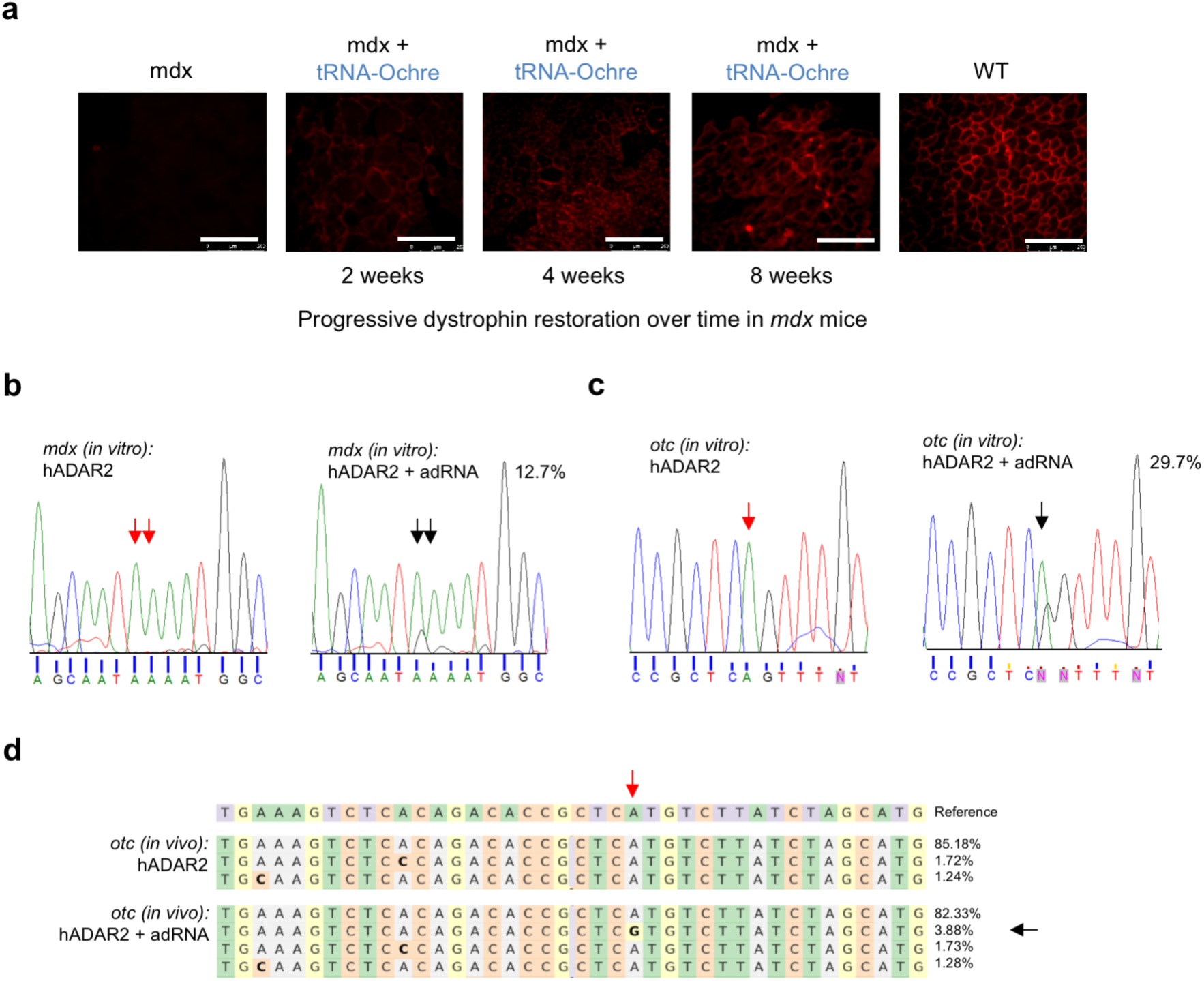
*In vitro* and *in vivo* targeting of DMD and OTC mRNA. **(a)** Progressively increasing restoration of dystrophin expression over time in *mdx* mice treated with AAV8-dual-serine-ochre-tRNA. **(b)** Representative Sanger sequencing plot showing 12.7% editing of the ochre stop codon in a fragment of the *mdx* dystrophin mRNA expressed in HEK 293T cells (quantified using NGS). **(c)** Representative Sanger sequencing plot showing 29.7% correction of the point mutation in a fragment of the *spf*^*ash*^ OTC mRNA expressed in HEK 293T cells (quantified using NGS). **(d)** Representative analyses of treated *spf*^*ash*^ mice showing 3.88% RNA correction of the point mutation (quantified using NGS).

